# A comprehensive evaluation of multiband-accelerated sequences and their effects on statistical outcome measures in fMRI

**DOI:** 10.1101/076307

**Authors:** Lysia Demetriou, Oliwia S Kowalczyk, Gabriella Tyson, Thomas Bello, Rexford D Newbould, Matthew B Wall

## Abstract

Accelerated functional Magnetic Resonance Imaging (fMRI) with ‘multiband’ sequences is now relatively widespread. These sequences can be used to dramatically reduce the repetition time (TR) and produce a time-series sampled at a higher temporal resolution. We tested the effects of higher temporal resolutions for fMRI on statistical outcome measures in a comprehensive manner on two different MRI scanner platforms. Experiment 1 tested a range of acceleration factors (1-6) against a standard EPI sequence on a single composite task that maps a number of basic sensory, motor, and cognitive networks. Experiment 2 compared the standard sequence with acceleration factors of 2 and 3 on both resting-state and two task paradigms (an N-back task, and faces/places task), with a number of different analysis approaches. Results from experiment 1 showed modest but relatively inconsistent effects of the higher sampling rate on statistical outcome measures. Experiment 2 showed strong benefits of the multiband sequences on results derived from resting-state data, but more varied effects on results from the task paradigms. Notably, the multiband sequences were superior when Multi-Voxel Pattern Analysis was used to interrogate the faces/places data, but showed less benefit in conventional General Linear Model analyses of the same data. In general, ROI-derived measures of statistical effects benefitted relatively little from higher sampling resolution, with decrements even seen in one task (N-back). Across both experiments, results from the two scanner platforms were broadly comparable. The statistical benefits of high temporal resolution fMRI with multiband sequences may therefore depend on a number of factors, including the nature of the investigation (resting-state vs. task-based), the experimental design, the particular statistical outcome measure, and the type of analysis used. Higher sampling rates in fMRI are not a panacea, and it is recommended that researchers use multiband acquisition sequences conservatively.

## Introduction

Acceleration in scanning speed is a long-standing goal of MRI research, and substantial gains in acquisition speed have been achieved by advances in both hardware and software. One major advance of particular interest to neuroimaging researchers is the development of ‘multiband’ or ‘Simultaneous Multi-Slice’ (SMS) sequences for functional MRI (Moeller et al., 2008). These use multiband excitation pulses to excite and collect multiple slices simultaneously, and provide increases in temporal resolution in line with the number of slices acquired at once; so a multiband factor of two acquires two slices simultaneously. This allows double the number of slices to be acquired in the same TR, or halves the repetition time (TR) needed for the same number of slices. High (up to 16) acceleration factors have been demonstrated (Moeller et al., 2008; 2010), that can substantially reduce the TR required for whole-brain imaging, and produce time-series with very high temporal resolution. However, as an undersampling technique, multiband sequences may produce decreased temporal signal to noise ratio (tSNR; Chen et al., 2015) and increased levels of images artefacts, in particular ‘slice-leakage’ effects (Barth et al., 2015; Todd et al., 2016). The trade-off between the benefit of higher temporal resolution and the cost of higher levels of noise and/or artefacts is important to characterize as these sequences become widely adopted.

The benefits of higher temporal resolution in fMRI may not be entirely obvious, considering that fMRI samples the BOLD (Blood-Oxygen-Level-Dependent) effect; a relatively low-frequency signal. Sampling this slow signal at a higher rate (beyond that necessary to adequately model it) may therefore seem to provide little benefit. However, BOLD effects are usually quantified using statistical methods, and those statistical tests depend crucially on the number of independent data points. Increasing the sampling rate reduces the influence of noise on statistical measures of the BOLD signal in much the same manner as more averaging of repeated measurements reduces the effect of noise and produces a more robust estimate (Miller, Bartsch, & Smith, 2016). Higher sampling rates can therefore potentially benefit the statistical outcome measures that researchers are often most interested in.

Previous work has shown that these sequences are indeed useful in this regard, within certain task domains or experimental approaches. Smith et al. (2010) used multiband sequences to increase the image resolution (2mm isotropic) of the entire brain with the same TR, and with signal-to-noise characteristics equivalent to standard EPI sequences. These sequences were used in the Human Connectome Project to generate high-resolution maps of functional connectivity using resting-state fMRI. Todd et al. (2016) recently evaluated multiband sequences at several acceleration factors (2, 4, and 6) and showed impressive gains on *t*-statistics, which varied depending on anatomical location, and the precise reconstruction algorithm used. Boyacioğlu et al. (2015) also demonstrated benefits of a Multi-Band Multi-Echo (MBME) sequence over a conventional multi-echo sequence at 7T, using both resting-state and task-activation data. Preibisch et al. (2015) found a substantial increase in sensitivity for resting-state analyses with four-fold acceleration, but also noted that higher acceleration levels produced artefacts.

While this previous work is useful, several unanswered questions remain. The majority of previous evaluations of multiband sequences have used resting-state fMRI data, with only a few using basic motor (finger-tapping) or visual (typically, gratings or checkerboards) stimulation paradigms (e.g. Todd et al., 2016; Boyacioğlu et al., 2015). These simple tasks are a classic method for evaluating fMRI sequences, but in many ways are quite dissimilar to the tasks used in modern cognitive neuroscience research, which may be relatively complex, and activate a wider network of brain regions than simple motor or sensory tasks. Secondly, there has been no published evaluation of the interaction between use of multiband-accelerated sequences and factors related to experimental design. Conceivably, higher temporal resolution scanning might be a particular benefit for fast event-related designs, relative to block designs. Thirdly, different analysis approaches have not been compared; the effect of multiband sequences on the signal-detection ability of conventional (i.e. General Linear Model-based) analysis of task data, relative to its effect on Multi-Voxel Pattern Analysis (MVPA) is one example that is currently undocumented. Finally, there have been no direct comparisons on the use of multiband sequences on different scanner platforms. Scanner hardware might reasonably be expected to have relatively minor effects, and a number of different scanner platforms have been used in previous evaluation work, but there has never been a direct comparison.

Our aim was therefore to address some of these questions, by performing as comprehensive a test of multiband acquisition sequences as possible, using several tasks, a number of different analysis approaches, and two different scanner platforms (a long, 60cm bore system, and a short 70cm bore system, both 3T). Our broad aim was to evaluate the ‘real-world’ performance of multiband sequences, using (currently) typical experimental and analysis techniques. We conducted two main experiments. The first sought to characterize the effect of a range of multiband acceleration factors (2-6) on a complex task that maps a number of sensory, motor, and cognitive networks. We then used a narrower range of acceleration factors (2 and 3) to comprehensively evaluate the statistical benefits of multiband sequences in three paradigms (two cognitive tasks, and resting-state data), with a number of different analysis approaches. We completed each experiment on both scanner platforms.

## Experiment 1 Methods

### Participants

Ten healthy volunteers were recruited for Experiment 1 of the study (5M, 5F, mean age = 24.6, range 20–39). Standard MRI screening procedures were followed for all participants in advance of testing. Informed consent was obtained from all the participants.

### Data Acquisition

Data was acquired on two scanners of the same field strength, but different RF, gradient, and magnet designs. Scanner 1 was a long bore 3T Siemens Tim Trio, and Scanner 2 was a short, wide bore 3T Siemens Magnetom Verio. The in-built body coil was used for RF excitation and the manufacturer’s 32 channel phased-array head coil was used for reception in both scanners. Whole-head anatomical images were acquired at the beginning of each scanning session using a Magnetization Prepared Rapid Gradient Echo (MPRAGE) sequence using parameters from the Alzheimer’s Disease Research Network (ADNI-GO; 160 slices × 240 × 256, TR = 2300 ms, TE = 2.98 ms, flip angle = 9°, 1 mm isotropic voxels, bandwidth = 240Hz/pixel, parallel imaging (PI) factor =2; Jack et al., 2008). B0 field images were acquired with a dual-echo gradient-echo sequence (TR = 599 ms, TE 1 = 5.19 ms, TE 2 = 7.65 ms, flip angle = 60°, 3 mm isotropic voxels, 35 axial slices, bandwidth = 260 Hz/pixel).

Six different functional sequences were used: a standard Echo-Planar Imaging (EPI) sequence, and five multiband acquisitions with different levels of acceleration: 1, 2, 3, 4 and 6 (hereafter referred to as MB1, MB2, MB3, MB4 and MB6). These sequences were based on the multiband EPI WIP v012b provided by the University of Minnesota (Auerbach et al., 2013; Cauley et al., 2014; Setsompop et al., 2012; Xu et al., 2013). Detailed characteristics of each sequence are shown in table 1. The sequences were standardized across the two scanners as much as possible, however because of the different hardware characteristics it was possible to set the bandwidth somewhat higher on Scanner 1 (2232 Hz/pixel) than on Scanner 2 (1906 Hz/pixel) due to gradient heating. A 3 mm isotropic resolution in a 192 mm FOV was acquired with interleaved slice acquisitions, and the TR was progressively shortened with increasing levels of multiband acceleration (down to a minimum of 333 ms for the MB6 sequence). The Ernst angle for each TR was used for excitation.

**Table 1.** Functional data acquisition sequences used in experiment 1.

| Scanner | Sequence | TR [ms] | Number of Slices | Number of volumes | Flip angle (degrees) | Echo time [ms] | Bandwidth [Hz/pixel] |
| --- | --- | --- | --- | --- | --- | --- | --- |
| 1 | Standard EPI | 2000 | 36 | 170 | 80 | 30 | 2232 |
|  | MB1 | 2000 | 36 | 170 | 80 | 30 | 2232 |
|  | MB2 | 1000 | 36 | 340 | 62 | 30 | 2232 |
|  | MB3 | 666 | 36 | 510 | 55 | 30 | 2232 |
|  | MB4 | 500 | 36 | 676 | 47 | 30 | 2232 |
|  | MB6 | 333 | 36 | 1006 | 40 | 30 | 2232 |
| 2 | Standard EPI | 2000 | 35 | 170 | 80 | 30 | 1906 |
|  | MB1 | 2000 | 35 | 170 | 80 | 30 | 1906 |
|  | MB2 | 1000 | 36 | 340 | 62 | 30 | 1906 |
|  | MB3 | 666 | 36 | 510 | 55 | 30 | 1906 |
|  | MB4 | 500 | 36 | 676 | 47 | 30 | 1906 |
|  | MB6 | 333 | 36 | 1006 | 40 | 30 | 1906 |

### Procedure and Tasks

Prior to the main experiment, a MRI phantom was used to collect one scan of each of the six sequences on each scanner. One hundred volumes of each sequence were collected, and these were used to calculate basic temporal signal-to-noise (tSNR) characteristics for all the sequences.

In the main experiment, participants viewed the visual stimuli through a mirror attached to the head coil that provided a view of a screen placed in the back of the scanner bore. Images were back-projected onto this screen through a waveguide in the wall of the scanner room. Auditory stimuli were delivered to the participant through MRI-compatible pneumatic headphones. Both scanners had similar audio-visual hardware.

The fMRI task (programmed using PsychoPy; Peirce, 2007) presented a battery of stimuli in order to assess an array of basic sensory and cognitive functions, and was adapted from Pinel et al.’s (2007) paradigm. The instructions/stimuli were either presented on screen (visual) or via the headphones (auditory). The four trial types were: a) visual gratings (high, medium, or low contrast; 30 trials), b) simple mental calculations (audio or visual instructions; 20 trials), c) pressing the left or right response key three times (visual or audio instructions; 20 trials) d) listening to or reading short sentences, e.g. “warm countries attract tourists” (20 trials). The combination of these four tasks and the variation in auditory and visual instructions allowed the mapping of five basic functional brain networks: visual, auditory, motor, cognitive, and language. The visual grating was a vertically-oriented sinewave patch with a Gaussian mask, which subtended approximately ten degrees of visual angle, had a period of 0.625 degrees of visual angle, and drifted to the right at a rate of 3.33 degrees/s. Three versions of the grating were generated varying in contrast level: high (100%), medium (25%) and low (5%). Randomly intermixed within the stimulus sequence were 20 null (blank screen) trials (also three seconds) in order to provide a baseline condition. Trials were presented in pseudo-randomised order in a single run of 110 trials of three seconds each.

Each complete scanning run lasted 5 minutes and 40 seconds. Seven versions of the battery task were created in which trials were presented in a different pseudorandom order. Participants performed the task seven times (one for each sequence: standard EPI, MB1, MB2, MB3, MB4, MB6, plus an additional standard EPI sequence; see analysis section below). The order of the acquisition sequences was randomised for each participant and each scanner, and subjects were blinded to which sequences were being performed during the scan. The total scanning session time was approximately 60 minutes. Participants completed two identical scanning sessions, one on each of the MRI scanners, also in a randomised order.

### Analysis

BOLD time-series from the 100-volume phantom scans were processed using custom MATLAB (Mathworks Ltd.) code and tSNR characteristics were calculated by dividing the temporal mean by the temporal standard deviation (Chen et al., 2015).

All the functional and anatomical data were preprocessed using FSL (FMRIB Software Library v5.0.4). BET was used for brain extraction of the anatomical data. Functional data preprocessing included motion correction, spatial smoothing with a 6 mm FWHM Gaussian process, high-pass temporal filtering (100 s), and a two-step coregistration to the subject’s individual anatomical image and an anatomical template image in standard stereotactic space (MNI152).

Data analysis for individual subjects was conducted in FSL’s FEAT module using the general linear model and FILM pre-whitening. Separate regressors were defined for the audio and visual variants of the motor, calculation, and language tasks, with three additional regressors modelling the three contrast levels of the visual grating stimulus, resulting in a total of nine task regressors. Head-motion parameters were included as regressors of no interest. The task regressors were convolved with a standard Gamma function (SD = 3 s, lag = 6 s) in order to model the HRF. Contrasts were computed that compared each individual task component with the baseline (null trials). For visual trials a mean contrast that compared the three grating conditions was compared to baseline.

The first-level analyses of all the subjects were combined into group level analyses using mixed effects (FLAME-1) models. A set of 14 group-level models were produced, one for each acquisition sequence performed on each scanner. A statistical threshold of *Z* = 2.3 (*p* < 0.05 cluster-corrected for multiple comparisons) was used for all group analyses.

The group level results from one of the standard EPI sequences on each scanner were used purely as functional localizers, to define Regions Of Interest (ROIs). This ensured that the ROI definition used entirely separate data and was unbiased. The ROIs were based on a set of key regions corresponding to the major activation clusters in the task: 1) primary visual areas in the occipital lobe, 2) primary auditory areas in the temporal lobe, 3) motor cortex (left hemisphere), 4) number/magnitude regions in the intraparietal sulcus (Fias et al., 2003; Shuman & Kanwisher, 2004; Bueti & Walsh, 2009), and 5) Wernicke’s and Broca’s area in the left hemisphere (see figure 1). Data from these ROIs were then extracted from the other six sets of results produced on each scanner.

**Figure 1.**
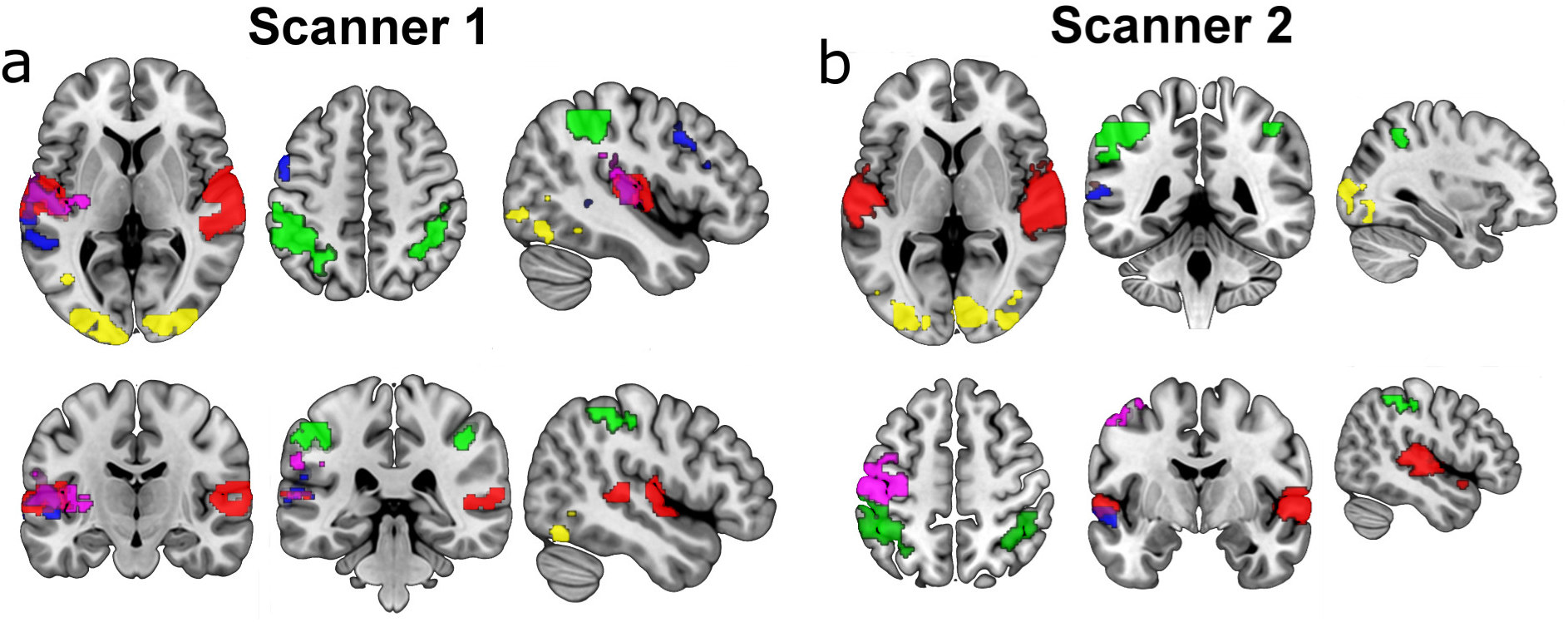
ROI masks used in experiment 1 (auditory: red, calculation: green, motor: pink, visual: yellow, language: blue) from scanner 1 (a) and scanner 2 (b). ROIs were defined based on an independent localiser scan conducted in the same session as the main experimental data, using a standard EPI sequence.

Two summary measures were calculated from the data. The first was a simple mean of the parameter estimate values within each ROI, re-calculated to represent % BOLD signal change from baseline; this reflects common practice in fMRI experiments. The second was the mean of the top 1% of Z scores across the brain, as previously used in Todd et al. (2016). This gives a measure of the top range of Z scores that is more reliable than simply using the peak score in the image. These two metrics were chosen as they relate directly to the amplitude of activation. Other measures that relate more to the spatial extent of activation clusters (such as number of activated voxels) may be problematic for multiband sequences because of ‘slice leakage’ effects, which reduce the independence between slices, can alias activations from one simultaneously-acquired slice to another, and create false positive activations at higher (4-6) acceleration levels. (Todd et al., 2016). Significant differences between both of these summary measures across the six differences were assessed using standard statistical methods (ANOVA and *t*-tests).

## Experiment 1 Results

### Temporal signal-to-noise measures

Figure 2a shows the results of the tSNR analysis performed in both scanners (Scanner 1 on the left and Scanner 2 on the right). There is an increase in tSNR for MB1 and MB2 compared to the Standard EPI, while MB3 is approximately the same level as the Standard EPI. Furthermore, a trend of reduced tSNR is observed in the more accelerated multiband sequences (MB4 and MB6) in both scanners.

**Figure 2.**
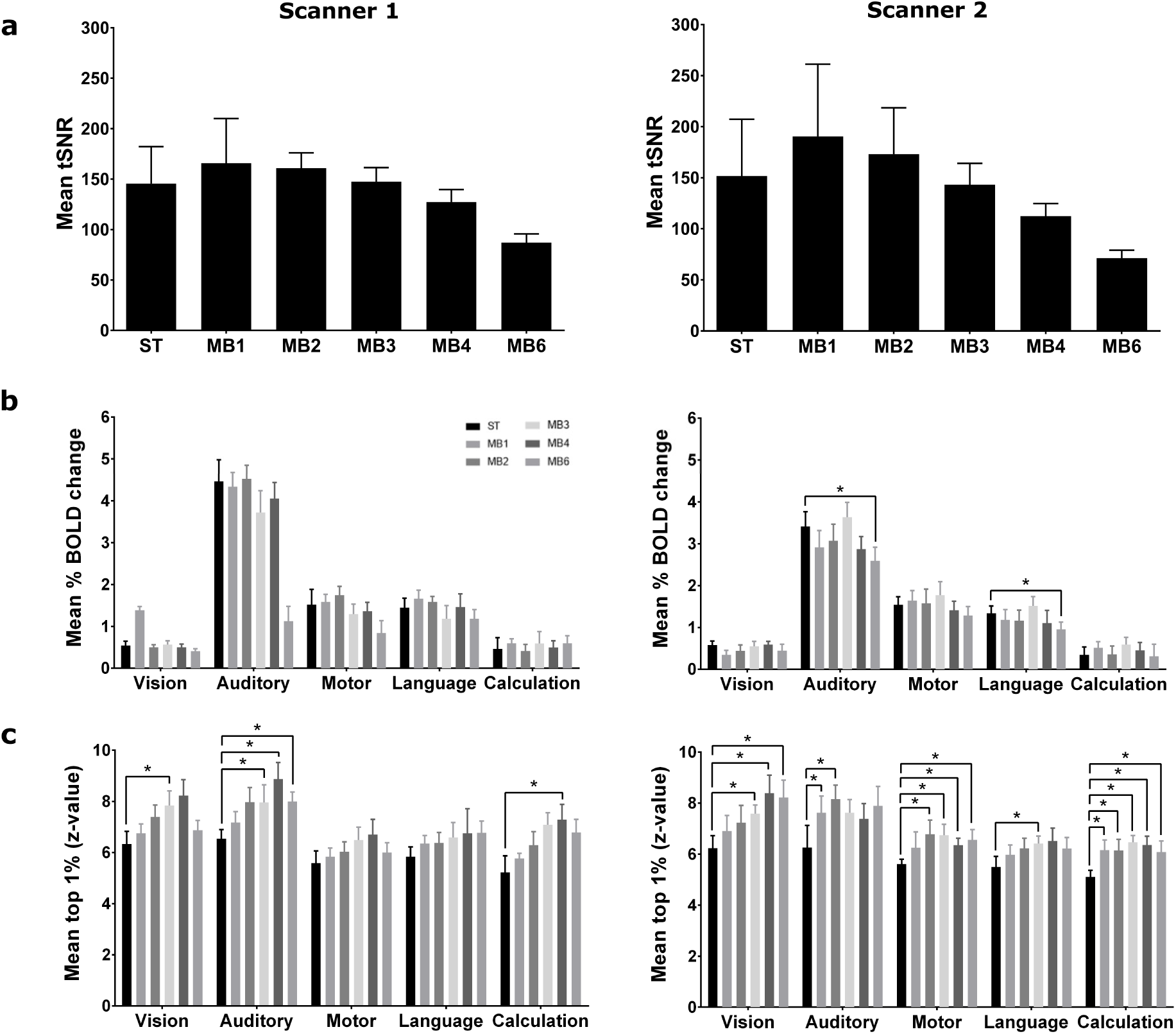
Results from experiment 1. a) Temporal signal-to-noise measures of the six sequences from both scanners. b) Mean % BOLD signal change from a set of independently-defined ROIs (* = *p* < 0.05). Only congruent ROI data is shown, i.e. vision columns show results from the visual task conditions, in the visual (occipital) ROI, auditory columns are the auditory task conditions in the auditory (superior temporal lobe) ROI, etc. c) Mean of the top 1% of *Z* scores in the statistical map from each contrast (* = *p* < 0.05). Error bars are standard errors.

### BOLD statistical maps

As expected, the whole-brain analysis for the fMRI battery task revealed significant activation in key areas across all the sequences tested in both scanners. However the strength and extent of activated voxels in each functional area varied (see figure 2). The standard EPI sequence produces adequate BOLD activation, with active voxels in the MB1, MB2, and MB3 sequences showing a similar pattern of intensity and spatial extent. However, the MB4 and MB6 maps are somewhat poorer, with reduced activation clusters for the motor and language tasks in particular. In addition visual activation in the occipital lobe is reduced in the fastest MB4 (Scanner 1) and MB6 (both scanners) sequences.

### ROI analysis

ROIs (see figure 1) were defined based on independent data collected during each scanning session. A 2 (Scanner) by 6 (acquisition sequence) by 5 (trial condition) ANOVA was performed on the ROI data, and showed a main effect of scanner (*F*[1,9] = 7.463, *p* = 0.023), a main effect of task condition (*F*[4,36] = 110.941, *p* < 0.001) and an interaction between scanner and task condition (*F*[4,36] = 15.177, *p* < 0.001). Since our primary interest is comparing the multiband sequences with the standard EPI sequence, *post hoc* analyses with *t*-tests focussed on this aspect of the data. These showed that the six sequences perform relatively comparably in terms of mean % BOLD signal within ROIs (see figure 2b). Sensitivity appears to drop off somewhat with the highest acceleration factors, and the reduced mean % BOLD signal in MB6 compared to the standard EPI is statistically significant for the auditory and language trials on Scanner 2. See table 2 for all statistical results.

**Table 2.** Paired *t*-test results of ROI data from experiment 1, comparing the standard EPI sequence with the multiband sequences, for all task conditions. All *p* values are two-tailed, and all degrees of freedom = 9. Significant (<0.05) *p* values are highlighted in bold text.

| Scanner | Condition | ST EPI - MB1 |  | ST EPI - MB2 |  | ST EPI - MB3 |  | ST EPI - MB3 |  | ST EPI - MB6 |  |
| --- | --- | --- | --- | --- | --- | --- | --- | --- | --- | --- | --- |
|  |  | *t* | *p* | *t* | *p* | *t* | *p* | *t* | *p* | *t* | *p* |
| 1 | Visual | -0.216 | 0.834 | 0.291 | 0.777 | -0.219 | 0.831 | 0.145 | 0.888 | 0.968 | 0.358 |
|  | Auditory | 0.246 | 0.811 | -0.136 | 0.895 | 0.961 | 0.362 | 0.581 | 0.575 | 1.155 | 0.278 |
|  | Motor | -0.168 | 0.870 | -0.642 | 0.537 | 0.575 | 0.580 | 0.350 | 0.734 | 1.958 | 0.082 |
|  | Language | -0.775 | 0.458 | -0.515 | 0.619 | 0.693 | 0.506 | -0.040 | 0.969 | 0.992 | 0.347 |
|  | Calculation | -0.671 | 0.519 | 0.213 | 0.836 | -0.344 | 0.738 | -0.106 | 0.918 | -0.592 | 0.568 |
| 2 | Visual | 2.122 | 0.063 | 0.810 | 0.439 | 0.220 | 0.831 | 0.449 | 0.664 | 0.951 | 0.366 |
|  | Auditory | 1.273 | 0.235 | 0.955 | 0.365 | -0.537 | 0.604 | 1.435 | 0.185 | 4.114 | **0.003** |
|  | Motor | -0.343 | 0.739 | -0.093 | 0.928 | -1.212 | 0.256 | 0.397 | 0.701 | -0.343 | 0.739 |
|  | Language | 0.595 | 0.566 | 0.691 | 0.507 | -0.567 | 0.585 | 0.734 | 0.481 | 3.386 | **0.008** |
|  | Calculation | -0.898 | 0.393 | -0.057 | 0.955 | -1.733 | 0.117 | -0.411 | 0.690 | 0.149 | 0.885 |

### Highest 1% of activated voxels

An ANOVA (with the same design as in the previous section) on these data showed no significant main effect of scanner (*F*[1,9] = 0.167, *p* = 0.692), but significant main effects of acquisition sequence (*F*[5,45] = 4.388, *p* = 0.002) and trial condition (*F*[4,36] = 10.883, *p* < 0.001). Also present were interactions between scanner and trial condition (*F*[4,36] = 3.696, *p* = 0.013), acquisition sequence and trial condition (*F*[20,180] = 1.817, *p* = 0.022) and all three factors (*F*[20,180] = 2.548, *p* < 0.001). *Post hoc* tests again focussed on the critical comparison between the standard and multiband sequences. Here the multiband-accelerated sequences generally out-performed the standard sequence. The mean of the top 1% of activated voxels in each of the five contrasts/ROIs showed a trend of increasing Z scores across all contrasts in both scanners (see figure 1c). The statistical results showed that the gains on Scanner 1 were marginal, with few significant differences. However, the increase in the top range of Z scores on Scanner 2 was more reliable, with a coherent pattern of significant increases in Z scores across multiband acceleration factors in the majority of the contrasts (excepting the auditory and language aspects of the task). See table 3 for all statistical results.

**Table 3.** Paired t-test results from experiment 1 of the highest 1% of activated voxels, comparing the standard EPI sequence with Multiband sequences. All *p* values are two-tailed, and all degrees of freedom = 9. Significant (<0.05) *p* values are highlighted in bold text.

| Scanner | Condition | ST EPI - MB1 |  | ST EPI - MB2 |  | ST EPI - MB3 |  | ST EPI - MB4 |  | ST EPI - MB6 |  |
| --- | --- | --- | --- | --- | --- | --- | --- | --- | --- | --- | --- |
|  |  | *t* | *p* | *t* | *p* | *t* | *p* | *t* | *p* | *t* | *p* |
| 1 | Visual | -0.764 | 0.465 | -1.178 | 0.269 | -2.364 | **0.042** | -1.887 | 0.092 | -0.758 | 0.468 |
|  | Auditory | -1.279 | 0.233 | -1.908 | 0.089 | -2.574 | **0.030** | -2.853 | **0.019** | -2.735 | **0.023** |
|  | Motor | -0.547 | 0.598 | -0.627 | 0.546 | -1.512 | 0.165 | -1.717 | 0.120 | -0.660 | 0.526 |
|  | Language | -1.789 | 0.107 | -0.782 | 0.455 | -1.119 | 0.292 | -0.950 | 0.367 | -1.548 | 0.156 |
|  | Calculation | -0.988 | 0.349 | -1.293 | 0.228 | -1.793 | 0.107 | -2.222 | 0.053 | -1.920 | 0.087 |
| 2 | Visual | -0.971 | 0.357 | -1.502 | 0.167 | -2.402 | **0.040** | -2.911 | **0.017** | -2.621 | **0.028** |
|  | Auditory | -0.827 | 0.430 | -3.697 | **0.005** | -2.217 | 0.054 | -1.540 | 0.158 | -1.916 | 0.088 |
|  | Motor | -1.235 | 0.248 | -2.201 | 0.055 | -2.406 | **0.040** | -2.312 | **0.046** | -2.525 | **0.032** |
|  | Language | -1.128 | 0.289 | -1.982 | 0.079 | -1.721 | 0.119 | -2.350 | **0.043** | -1.859 | 0.096 |
|  | Calculation | -2.687 | **0.025** | -2.679 | **0.025** | -4.651 | **0.001** | -5.550 | **0.001** | -2.274 | **0.049** |

**Figure 3.**
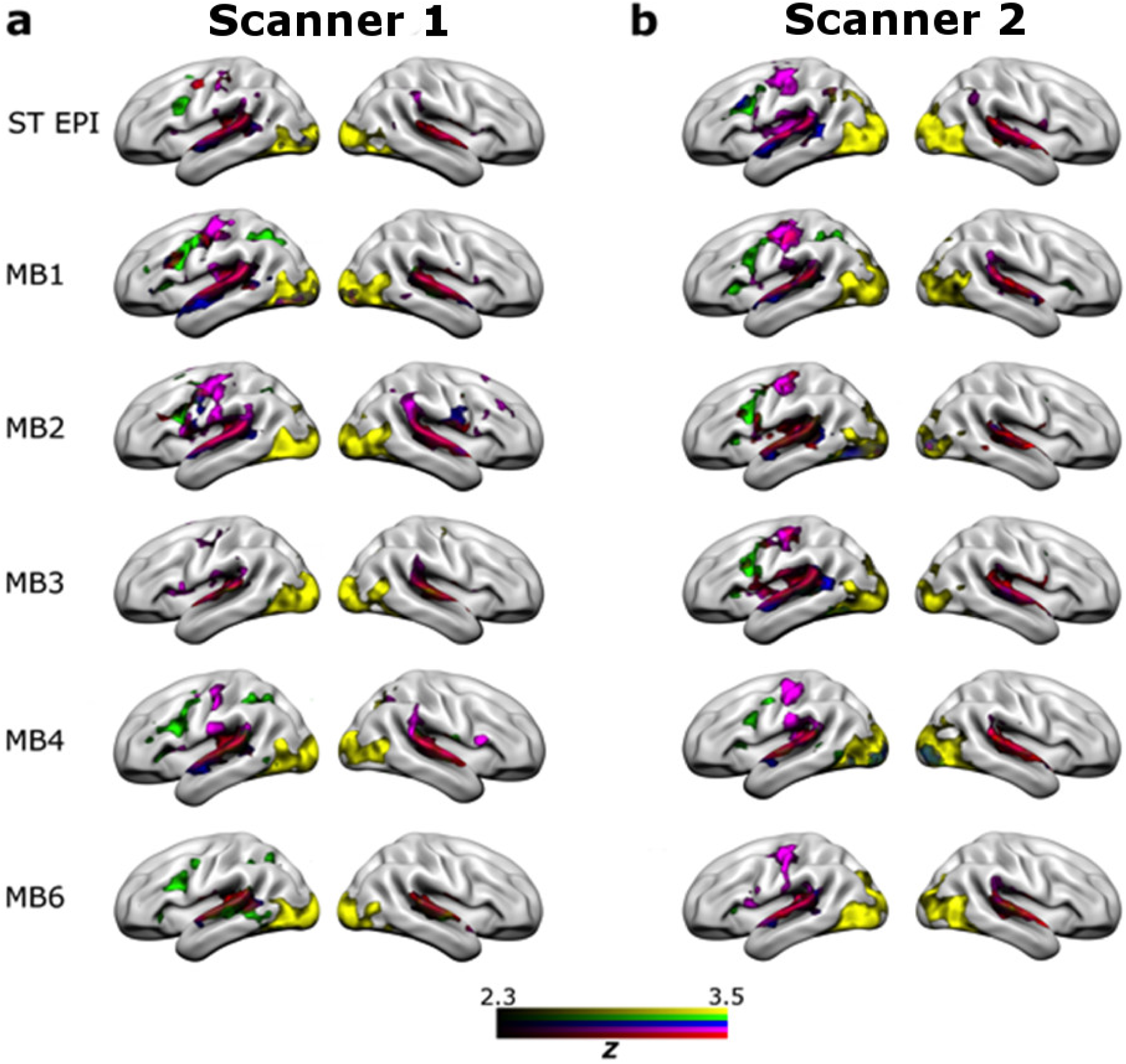
Group-level statistical maps for each MRI sequence (Standard EPI, MB1, MB2, MB3, MB4, MB6) showing results for the five task conditions assessed in the fMRI battery task: vision (yellow), auditory (red), motor (pink), calculations (green) and language (blue). Results from Scanner 1 are on the left and results from Scanner 2 are on the right. All statistical maps are thresholded at *Z* = 2.3, *p* < 0.05 (cluster corrected for multiple comparisons).

## Experiment 1 Discussion

Results from experiment 1 were somewhat mixed, with a clear decrease in tSNR at higher acceleration factors, and only marginal differences when conventional analysis methods (calculating % BOLD signal change in ROIs) are used. However, the analysis of the mean of the top 1% of *Z* scores showed some benefit of the multiband sequences on the top range of the statistical results, suggesting somewhat stronger effects and more robust statistics.

The task used in experiment 1 was a fast event-related paradigm, designed to map a number of basic sensory, motor, and cognitive functions in as short a time as possible. This task was chosen as it provides several different readouts, and its short duration still allowed seven repetitions in a single scanning session without excessive subject fatigue. However, the design is not entirely typical for an fMRI experiment, with short trials, presented almost continuously. It is possible that the statistical benefit of short-TR multiband sequences might be more evident with a different task design. Taking into consideration all the results of experiment 1, the highest-performing sequences (in terms of tSNR, and the results from the main experiment) were MB2 and MB3. It was therefore decided to test the MB2 and MB3 sequences against the standard EPI sequence, on several different tasks, and using a variety of analysis approaches. This was the aim of experiment 2.

## Experiment 2 Methods

### Participants

Fourteen healthy volunteer participants were recruited and tested on each scanner (Scanner 1: 7M, 7F, mean age = 24.86 range = 21-33; Scanner 2: 9M, 5F, mean age = 26.36, range = 21-39). Standard MRI screening procedures were followed for all participants in advance of testing. Informed consent was obtained from all the participants.

### Data Acquisition

The standard EPI, MB2, and MB3 sequences used in experiment 2 were the same as those used in experiment 1 (see table 1), with the only difference being the number of volumes acquired in each sequence (see description of tasks below). High-resolution T1 images and B0 field-maps were also acquired at the beginning of each session, also using the same sequences as experiment 1 and described above. As in experiment 1, data was acquired on both scanner platforms.

### Procedure and Tasks

Experiment 2 employed a within-subjects design with an event-related design task, a block-design task, and a resting state scan. Both the tasks were programmed in PsychoPy (Peirce, 2007).

The event-related paradigm was a faces/places task (Epstein & Kanwisher, 1998; O’Craven & Kanwisher, 2000; Pegors et al., 2015). Face images were taken from the Chicago Face Database (Ma, Correll, & Wittenbrink, 2015), and were balanced for gender and ethnicity. Happy and fearful facial expressions from the same 12 individual were used, with 24 stimuli in total. The ‘places’ stimuli were acquired through internet searches of standard stock image libraries using Google image search (all royalty/copyright free images, labelled for reuse). Twelve ‘positive’ place images (attractive neighbourhoods, peaceful landscapes, etc.) and 12 ‘negative’ images (war-torn landscapes, bombed buildings, etc.) were acquired and used in the task. Each image was presented for 2 seconds, during which the participants were asked to classify each image as either “positive” or “negative” using two keys (index and middle finger, respectively) on a response box. Presented at the bottom of the screen was a small schematic of a hand with the responses marked near the index and middle fingers, as a reminder of the response mappings. Inter-trial-intervals (ITIs) of variable duration (2-10s) with a mean of 5.5s and an approximately Poisson distribution (Hagberg et al., 2001) were used, during which the screen was blank. There were 48 trials, and three different pseudo-random stimulus sequences were programmed for use in the three repetitions of the task in each scanning session. The total duration of the task was 6 minutes.

The block-design paradigm was an N-back task designed to tax working memory capacity, adapted from Ragland et al (2002). Alternating 0-back and 2-back blocks were presented. For the 0-back blocks, the participants had to remember an initial target letter and respond whether the subsequent letters matched the target. In the 2-back blocks, the participants had to recall whether each letter presented on the screen matched the letter that was presented two trials before. Participants responded using an MRI-compatible response box, with the index and middle finger of the right hand used for ‘yes’ and ‘no’ responses, respectively. Each block lasted 20 seconds, contained 10 two-second trials, and was followed by a 10 second rest period. Six repetitions of each block type were presented, for a total task time of six minutes.

The final paradigm was a six minute resting state scan. Participants were instructed to keep their eyes open, and to relax. A blank screen was displayed for the duration of the scan.

The first scan was always the faces/places task, followed by the N-back task, followed by the resting-state, and this order was maintained throughout the scanning session. Each task was presented three times for a total of nine in a session, with a different acquisition sequence (standard EPI, MB2, or MB3) used for each set of three tasks. The order of acquisition sequences was randomised across participants, and all participants were blinded to which acquisition sequence was being used on which tasks while in the scanner. Scanning sessions on each scanner were identical, and 12 of the 14 subjects completed both sessions. Four subjects completed sessions on only one scanner.

### Analysis

Pre-processing of the anatomical and functional data was exactly the same as in experiment 1.

Analysis of the N-back task in FSL’s FEAT module used the 0-back and 2-back blocks modelled as explanatory variables, and also included standard head-motion regressors. The same Gamma function was used to model the HRF as in experiment 1. Contrasts were computed to model the effects of each condition alone (relative to the baseline segments of the time-series), and 2-back > 0-back.

For the faces/places task the experimental conditions were: face +ve, face −ve, place +ve, and place −ve. Individual regressors were produced for each, and the model also included the first temporal derivative of each regressor, as well as the standard head-motion parameters. The HRF was modelled with the same standard Gamma function. Contrasts were computed to model the effects of each condition alone (relative to baseline) as well as to examine the effects of faces > places (and vice versa) and positive > negative (and vice versa).

For the resting-state data, seed-based connectivity analyses were performed using Posterior Cingulate Cortex (PCC) and anterior insula masks (see figure 4), both derived from Neurosynth (using thresholded versions of the ‘default’ and ‘salience’ term maps, respectively; http://neurosynth.org/analyses/terms/default/ and http://neurosynth.org/analyses/terms/salience/). These standard-space masks were back-projected into individual subject space, and time-series were extracted from the ROIs for each individual subject. These formed the basis for defining three networks, the Default-Mode Network (DMN; defined as positive connectivity with the PCC seed region; Fox & Raichle, 2005), the Executive Control Network (ECN, defined as negative connectivity with the PCC seed region; ibid.) and the Salience Network (defined as positive connectivity with the anterior insula seed-region; Seeley et al., 2007; Goulden et al., 2014). Individual subjects’ anatomical images were segmented (using FSL’s FAST module), and mean white matter and CSF time-series were produced, using the separate anatomical masks. These were included in the model as regressors of no interest, along with standard head-motion regressors.

**Figure 4.**
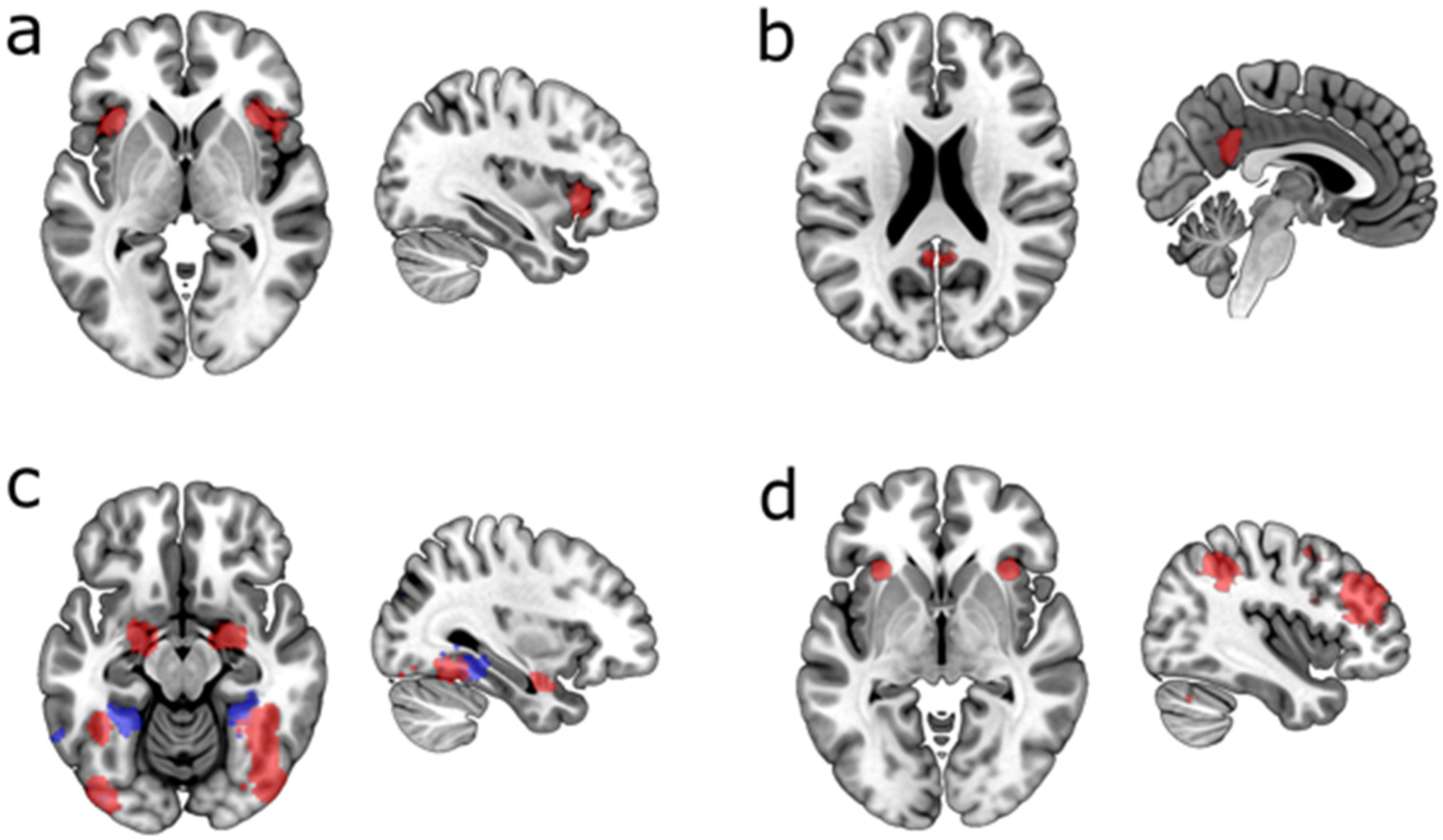
ROI masks for tasks of Experiment 2. a) Anterior Insula seed-region used to define the salience network. b) PCC seed-region, used to define the default mode and executive control networks. c) Faces/Places mask (red: face, blue: place). d) Working memory ROI mask used in the N-back task. All masks derived from statistical maps downloaded from http://www.neurosynth.org/ (see main text), in order to provide ROI definitions independently from the main experimental data.

The first-level analyses of all the paradigms were combined into group analyses using mixed effects (FLAME-1) models. As for experiment 1, a statistical threshold of *Z* = 2.3, (*p* < 0.05 cluster-corrected for multiple comparisons) was used for all group analyses. A set of six group-level analyses were produced for the N-back and faces/places task (three sequences, tested on the two scanners) and a set of 18 models was produced for the resting-state data (three different defined networks, using three sequences, on two scanners). Subsequent ROI analysis was conducted using ROIs derived from the ‘working memory’ term on Neurosynth (http://neurosynth.org/analyses/terms/working%20memory/) for the N-back task. ROIs for the faces/places task were defined based on the ‘faces’ and ‘place’ terms on Neurosynth (http://neurosynth.org/analyses/terms/faces/ and http://neurosynth.org/analyses/terms/place/). This independent definition of the ROIs based on automated meta-analysis of the previous literature provided an unbiased and objective ROI definition. Data was extracted from these ROIs for each task condition in both tasks, relative to baseline, for each sequence, and for data from each scanner. As in experiment 1, two different summary measures were computed; a ROI-based measure of BOLD percentage signal change from baseline, and a measure of the top 1% of *Z* values in the statistical maps. The latter measure was also used to quantify the resting-state data.

Additional analyses of the resting-state data used the dual regression procedure (Beckmann et al., 2009) and a set of 10 canonical resting-state networks identified by Smith et al. (2009). Data were pre-processed using FSL’s Melodic module with the same settings for pre-processing as used previously, and dual regression was carried out using the 10 Smith et al. (2009) networks as the inputs. Six separate dual regressions were carried out, one for each sequence on each scanner. This produced a set of 10 individualised networks for each subject/sequence/scanner, which could be compared using the top 1% of *Z* values metric.

Additional analyses were conducted on the faces/places task data using a Multi-Voxel Pattern Analysis (MVPA) approach, as instantiated in the Pattern Recognition of Brain Image Data (PROBID, http://www.brainmap.co.uk/probid.htm, version 1.04) toolbox for Matlab. This approach was used to test the classification of the positive/negative dimension of the stimuli (as in Pegors et al., 2015), which we hypothesized to be weaker and less spatially distinct than the relatively well-replicated and stronger difference between face and place stimuli (e.g. Epstein & Kanwisher, 1998; O’Craven & Kanwisher, 2000). Both Support-Vector Machine (SVM) and Gaussian-Process Classifiers (GPC) were used to separately test the three sequences from the two scanners, with a whole-brain mask used for all analyses. Inputs to the classification algorithms were the contrast files (positive > baseline, and negative > baseline) resulting from the first-level univariate analyses. Permutation tests were used to determine the statistical significance of the conducted analyses. For each comparison *p*-values were obtained by permuting the class labels 1000 times.

## Experiment 2 Results

### Faces/Places Task

Inspection of the statistical maps from the three sequences (see figure 8a) showed a broadly similar pattern of results, with hippocampal and para-hippocampal regions responding strongly to place stimuli and ventral visual regions in the fusiform responding preferentially to face stimuli (in line with previous work, e.g. Epstein & Kanwisher, 1998). ROI data was analysed using a 2 (scanner) by 3 (acquisition sequence) by 4 (task condition) ANOVA model. This showed only a significant main effects of task condition (*F*[3,39] = 22.17, *p* < 0.001). Hypothesis-driven *post hoc* tests again compared multiband to the standard sequence. A significant difference is only seen between the standard EPI and MB3 sequences for the face −ve stimuli on Scanner 1 (see table 4, and figure 5a). Analysis of the highest 1% of activated voxels using the same ANOVA model showed significant main effects of acquisition sequence (*F*[2,26] = 28.79, *p* < 0.001) and task condition (*F*[3,39] = 67.537, *p* < 0.001). *Post hoc* comparisons revealed a consistent significant increase for both multiband sequences compared to the standard EPI, across all stimuli and both scanners (excepting place −ve stimuli on Scanner 2; figure 5b and table 5). Differences between MB2 and MB3 were small, and not statistically robust.

**Figure 5.**
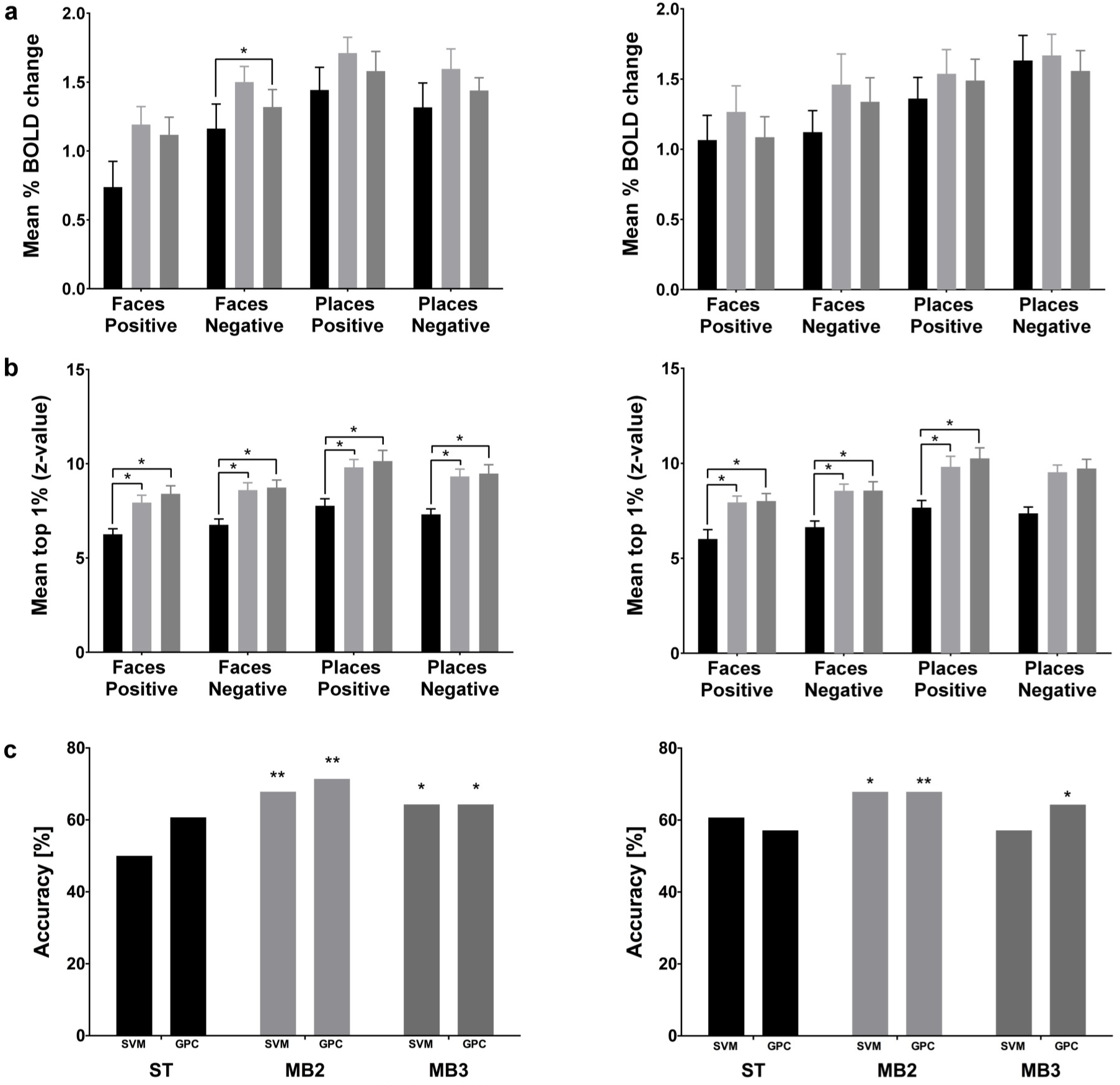
Results for Faces/Places task in Experiment 2 comparing the Standard EPI sequence with MB2 and MB3. a) Mean % BOLD signal change from the ROI masks for faces and places (see figure 4). b) Mean of the top 1% of *Z* scores in the statistical map from the faces/places contrasts c) MVPA analysis % accuracy results for each sequence using both Support Vector Machine (SVM) and Gaussian Process Classifier (GPC) algorithms. * = *p* < 0.05, ** = *p* < 0.01. Error bars are standard errors.

Further analysis on the valence dimension of the stimuli using MVPA showed poor performance (not significantly different to chance/50%) of the classifier algorithms for the standard EPI sequence (figure 5c). However, classification performance on the MB2 and MB3 sequences was improved, and statistically reliable (with the exception of the MB3 sequence using the SVM classifier, on Scanner 2). This suggests that the MB2 and MB3 sequences are better at providing distinctive patterns of BOLD activation between positive and negative images.

### N-back Task

The patterns of BOLD activation found in this task were again reasonably similar across the three sequences (see figure 8b) and also consonant with previous work using similar tasks (Owen et al., 2005). A 2 (scanner) by 3 (acquisition sequence) by 2 (task condition) ANOVA showed only a main effect of task condition (*F*[1,13] = 181.809, *p* < 0.001). Comparison of sequences using *t*-tests revealed that mean % BOLD change (Figure 6a) is decreased across stimuli/tasks in the multiband sequences compared to the standard-EPI sequence. Specifically, MB2 on Scanner 1 shows a significant decrease for the 0-back, while MB3 shows significant decreases in both 0- and 2-back conditions. In contrast, analysis of the mean of the top 1% of voxels (Figure 6b), using the same ANOVA model showed significant main effects of task condition (F[1,13] = 5.841, p = 0.031) and acquisition sequence (*F*[2,26] = 49.311, *p* < 0.001). Detailed comparison reveals that MB2 and MB3 very consistently produced a significant increase in the top range of *Z* values in both task conditions (0- and 2-back), in both scanners (figure 6b and table 5).

**Figure 6.**
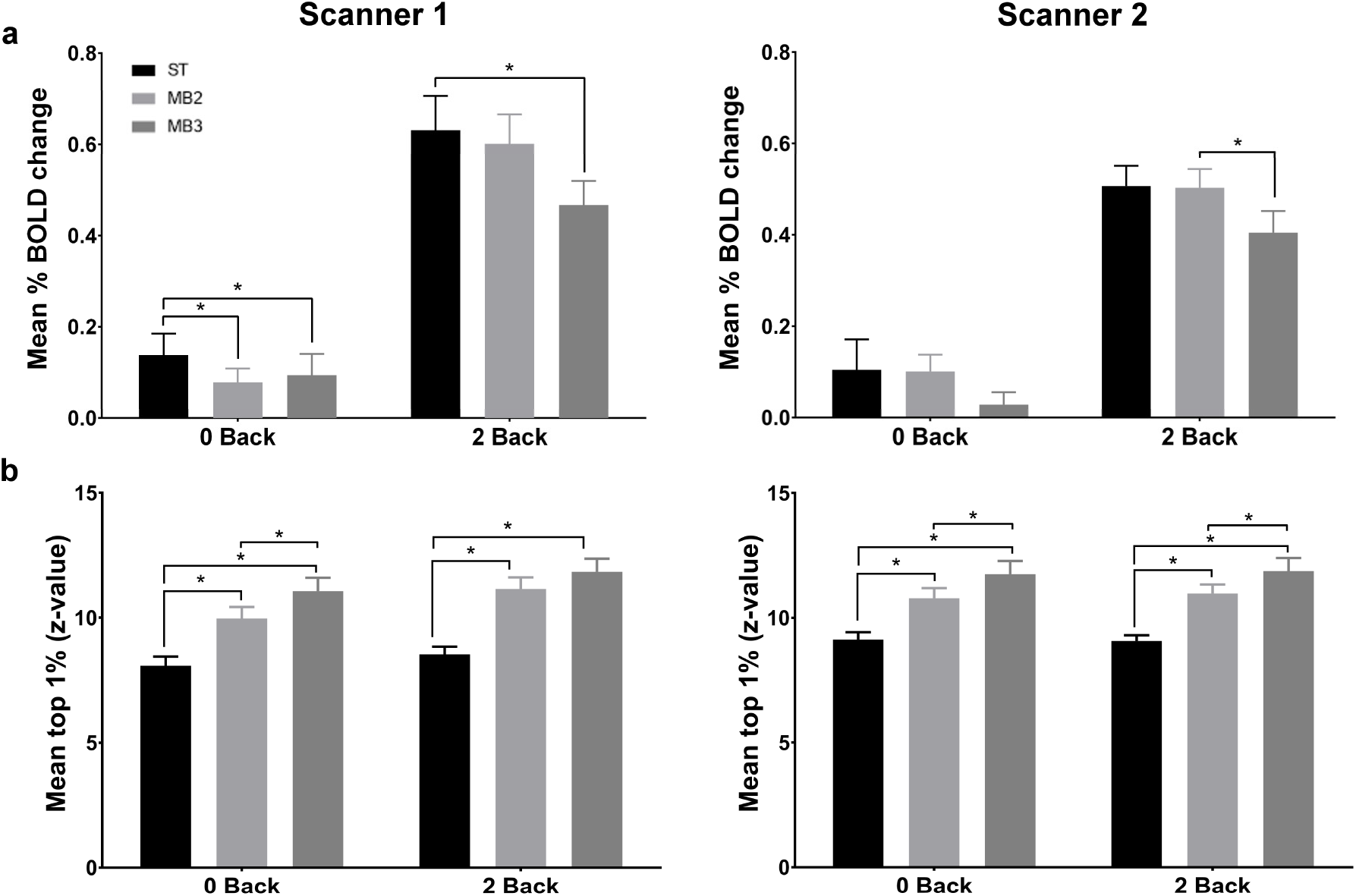
Results for N-back task in Experiment 2 comparing the Standard EPI sequence with MB2 and MB3. a) Mean % BOLD signal change from the ROI mask for working memory (see figure 4). b) Mean of the top 1% of Z scores in the statistical map from 0-back and 2-back contrasts. * = *p* < 0.05. Error bars are standard errors.

### Resting State Data

A statistical analysis was performed on the mean of the top 1% of activated voxels from the seed-based analysis of the three pre-determined resting state networks (DMN, ECN, and Salience network). A 2 (scanner) by 3 (acquisition sequence) by 3 (resting state network) ANOVA showed significant main effects of acquisition sequence (*F*[2,26] = 19.77, *p* < 0.001) and network (*F*[2,26] 830.311, *p* < 0.001), as well as an interaction between sequence and network (*F*[4,52] = 8.652, *p* < 0.001). Detailed comparisons show an increase in activation in MB2 for the Default Mode Network on both scanners (Figure 7a) and the Executive Control Network on Scanner 1. In addition MB3 shows increased activation in the Default Mode network on Scanner 2, and in the Salience Network on both scanners.

**Figure 7.**
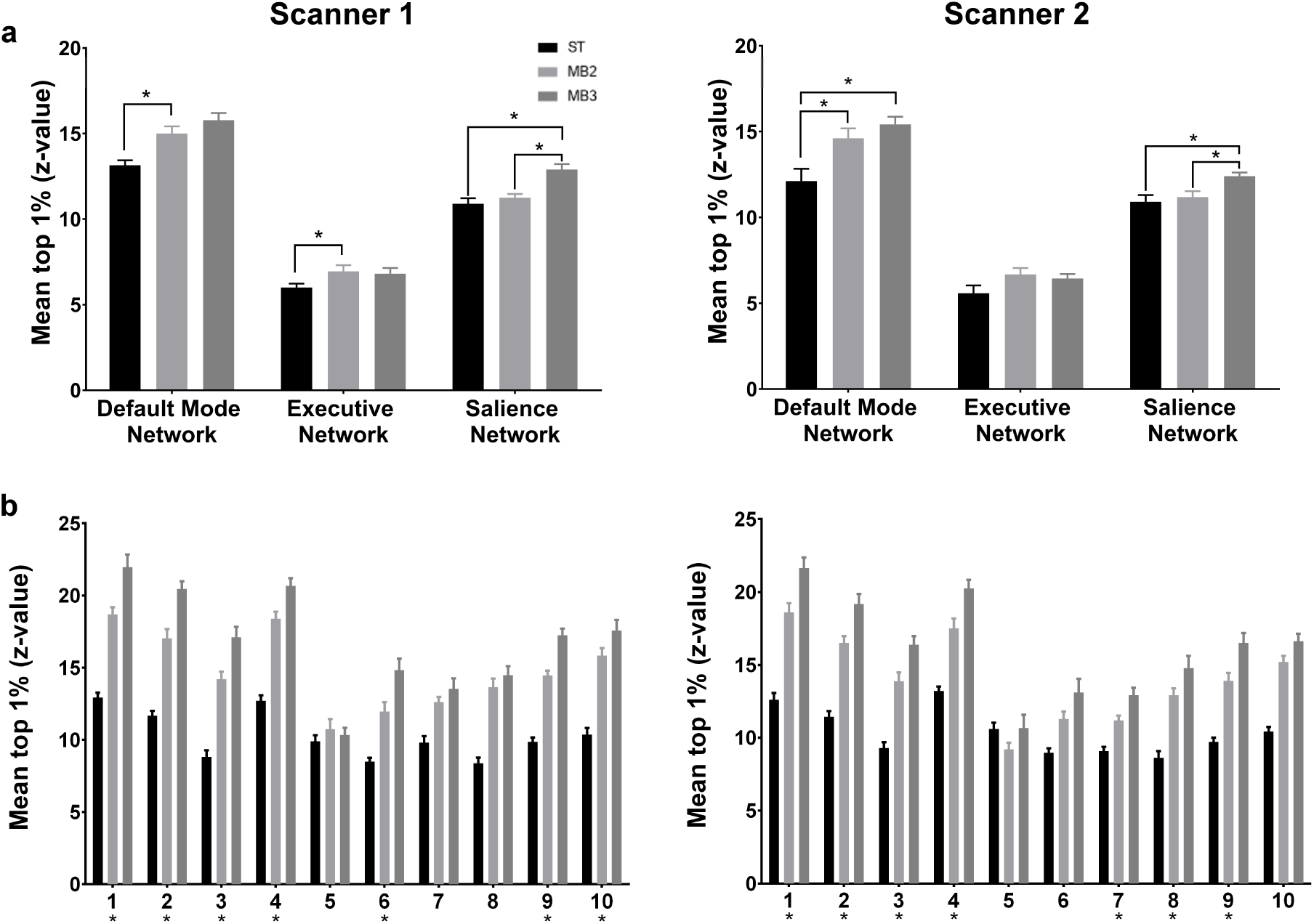
Results for Resting State data in Experiment 2 comparing the Standard EPI sequence with MB2 and MB3. Mean of the top 1% of Z scores in: a) three seed-region based analyses: PCC for Default Mode Network and Executive Control Network, and anterior insula for the Salience Network (see figure 4), * p < 0.05, and b) on the 10 resting state networks defined by Smith et al (2009), * (beneath the × axis) = *p* < 0.05 for all pairwise comparisons (i.e. ST-EPI vs. MB2, ST-EPI vs. MB3, MB2 vs. MB3). Error bars are standard errors.

A 2 (scanner) by 3 (acquisition sequence) by 10 (resting state network) ANOVA was performed on the 10 networks derived from Smith et al (2009). This showed significant main effects of acquisition sequence (*F*[2,26] = 419.938, *p* < 0.001) and network (*F*[2,26] 85.196, *p* < 0.001), as well as an interaction between sequence and network (*F*[4,52] = 20.399, *p* < 0.001). *Post hoc* tests showed a significant increase in the mean of the top 1% of Z values in the visual networks (1-3), the default mode network (4) and the left-lateralised fronto-parietal network (9) in both MB2 and MB3, and on both scanners. Furthermore, the multiband sequences produced higher statistical values in sensorimotor (6), and right-lateralised fronto-parietal (10) networks on Scanner 1, and in the auditory (7) and executive (8) networks on Scanner 2 (see figure 7b).

**Table 4.**
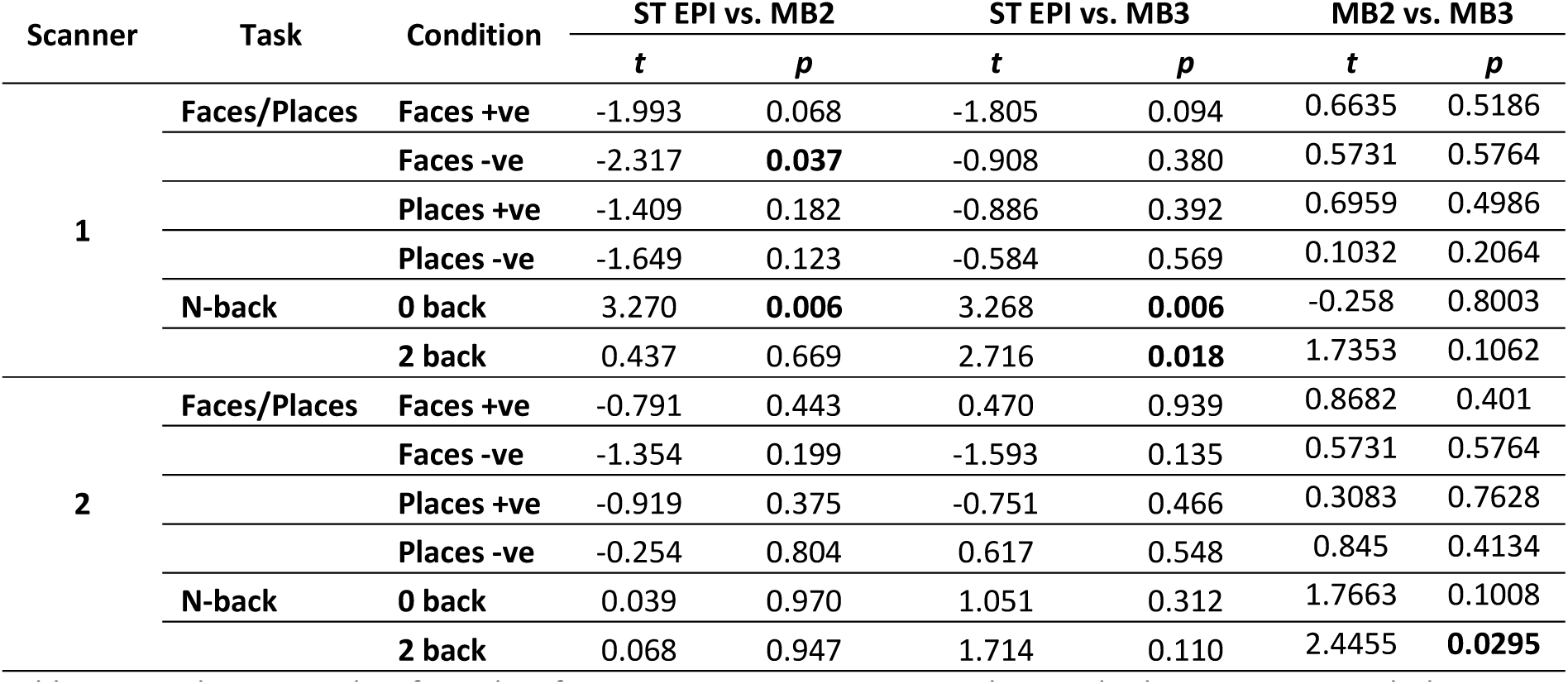
Paired *t*-test results of ROI data from experiment 2, comparing the standard EPI sequence with the MB2 and MB3 sequences. Significant (< 0.05) *p* values are highlighted in bold text. All *p* values are two-tailed, and all degrees of freedom = 13.

**Table 5.**
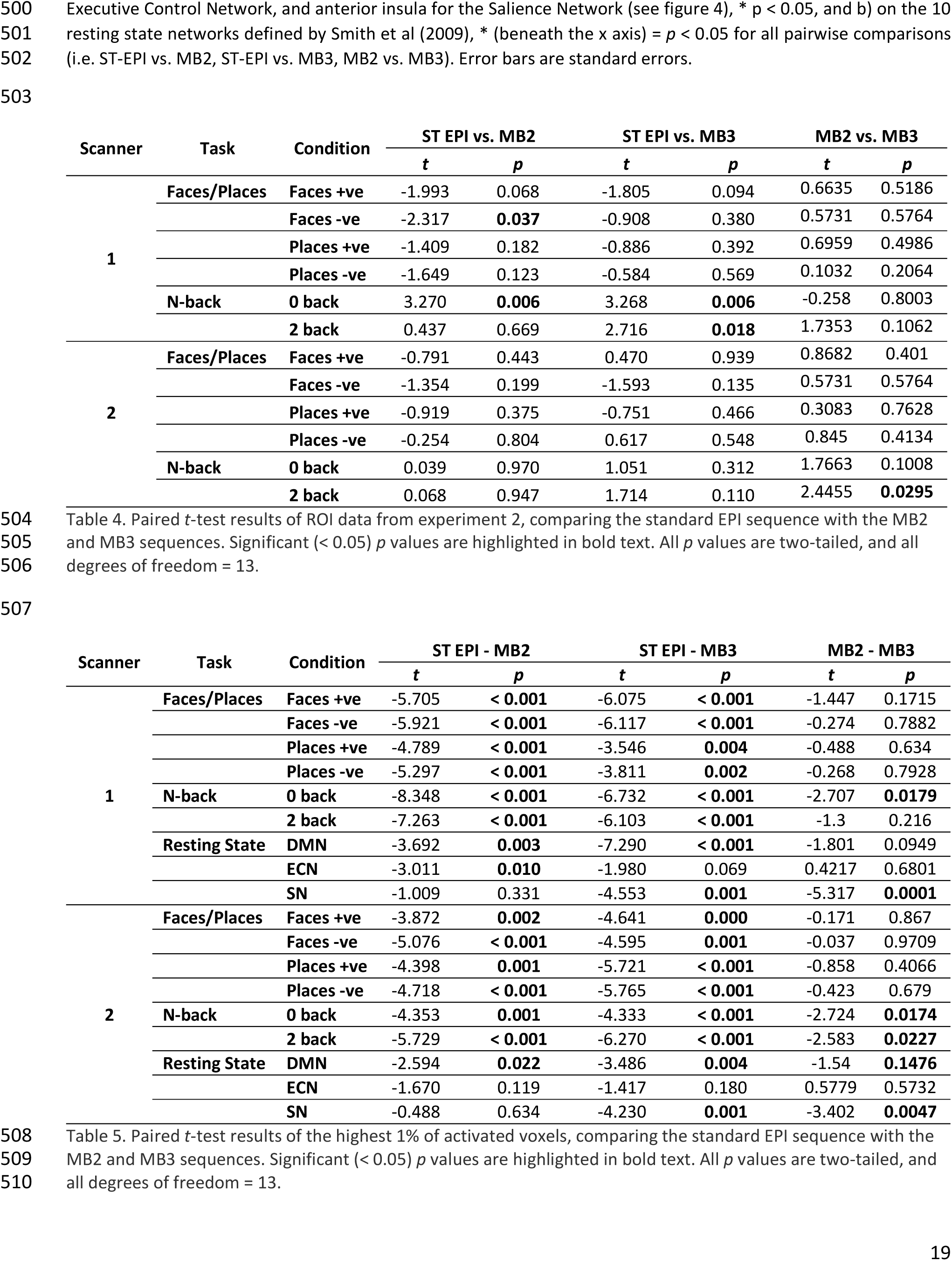
Paired *t*-test results of the highest 1% of activated voxels, comparing the standard EPI sequence with the MB2 and MB3 sequences. Significant (< 0.05) *p* values are highlighted in bold text. All *p* values are two-tailed, and all degrees of freedom = 13.

**Figure 8.**
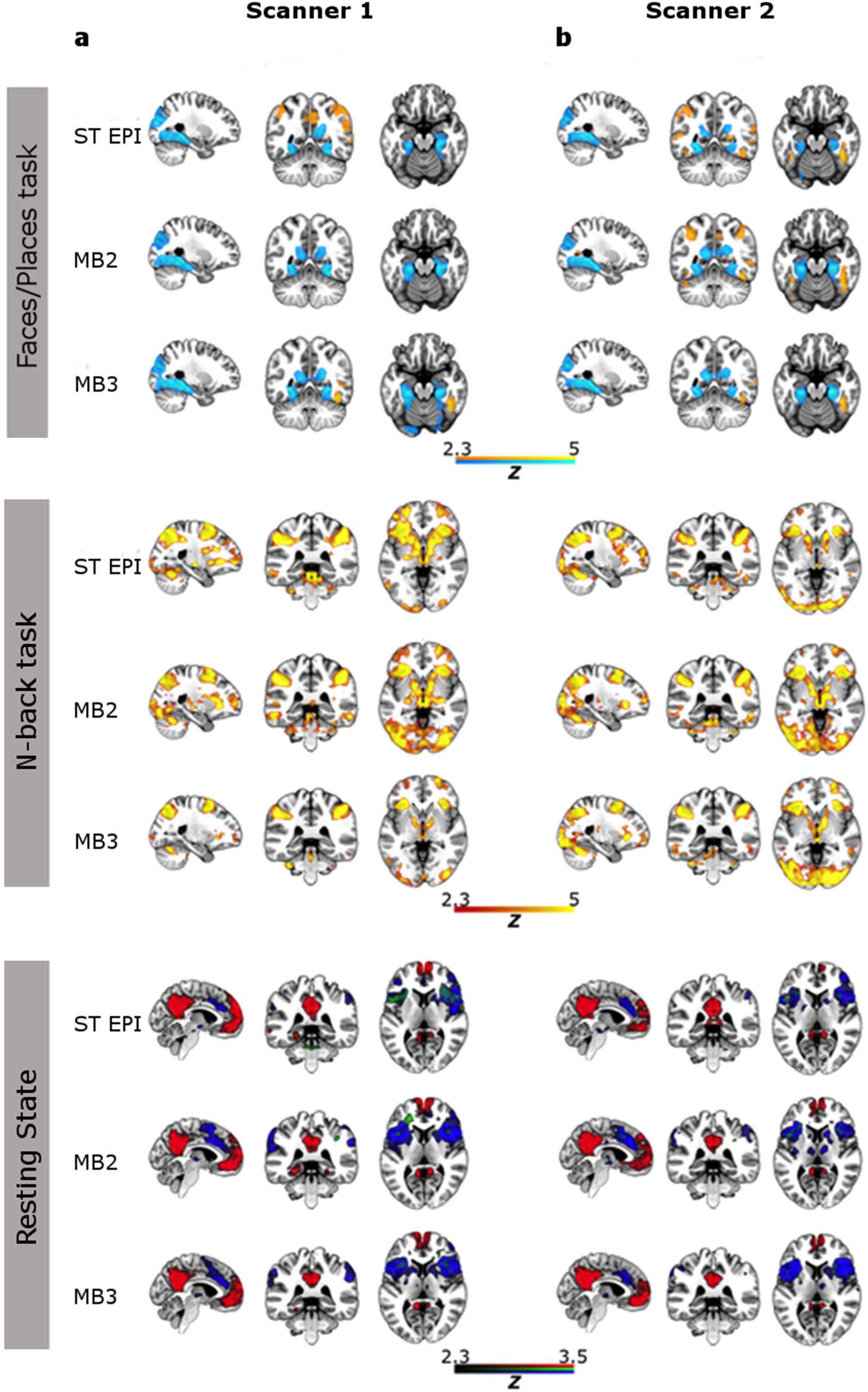
Group-level statistical maps for each MRI sequence (Standard EPI, MB2, MB3) in the three experimental paradigms used in experiment 2. a) Faces/Places task: results from the faces>places contrast in red-yellow and results from the places>faces task in blue-cyan. b) N-back task: results from the 2>0-back contrast in red-yellow. c) Resting State scan using the PCC and anterior insula seed-regions: Default Mode Network in red, Executive Control Network in green and Salience Network in blue. All statistical maps are thresholded at *Z* = 2.3, *p* < 0.05 (cluster corrected for multiple comparisons).

### General Discussion

Experiment 1 demonstrated that use of multiband sequences significantly increased the mean of the top range of statistical values in the results, though gains were relatively modest and not completely consistent across task conditions and the two scanners tested. Importantly, when conventional analysis techniques were used (calculating the mean of independently-defined ROIs), no particular advantage of higher temporal resolution scanning was evident. Experiment 2 showed much more consistent and substantive effects for both the faces/places and N-back tasks on the top range of statistical scores, but again, these benefits did not translate to more typical ROI-based analysis techniques, with significant *decreases* in measures of percentage signal change seen for the multiband sequences on the N-back task. The MVPA of the valence dimension of the faces/places task showed a strong benefit, with significant classification performance only seen in the MB2 and MB3 sequences. In the resting-state data from experiment 2, multiband sequences produced relatively modest (but consistent) increases in the top range of *Z* scores using seed-based analyses of the DMN, ECN and Salience network. However, impressive gains in sensitivity were seen in the resting-state data when an ICA-based dual regression (Beckmann et al., 2009) approach was used, with dramatic increases in the top range of statistical values across the majority of a set of 10 standard resting-state networks (derived from Smith et al., 2009).

Perhaps the most interesting aspect of these data is the discrepancy between a commonly-used measure (the mean of all voxels within a pre-defined ROI, expressed in % BOLD signal change) and a measure of the top range of statistical values (mean of the top 1% of Z scores; Todd et al., 2016). In the task data from experiments 1 and 2 the use of multiband sequences produced more reliable increases in the latter than the former. One possible explanation is that the multiband sequences may produce clusters with higher statistical reliability, but a smaller spatial extent. The statistical values within these small clusters may be (relatively) high, but when averaged in with background-level voxels in (larger) ROIs, their values become diluted. Another is that multiband sequences were created (and have been used most often) for increasing spatial resolution in EPI scanning, while maintaining relatively standard TR times (e.g. Smith et al., 2013). Smaller voxel size reduces susceptibility-related dropouts (Moeller et al., 2008) and better localizes the BOLD signal to the grey matter. The relatively large ROIs defined here probably include both grey and white matter voxels, which may further reduce the strength of the ROI results. This discrepancy between the two outcome measures highlights an issue with the use of mean ROI values; arguably a crude outcome measure, but still a widely-used one, despite clear demonstrations in the literature of more reliable and sensitive alternatives (e.g. Mitsis et al., 2008).

The multiband sequences even produced statistically reliable *decreases* in sensitivity in the N-back task, when using ROI measures of percentage signal change. This highlights the potential role of experimental design; in this block-design task, the advantage of averaging more data points per block may be marginal, as each block already contains a large number of TRs. Increases in leakage-factor related noise from the multiband acceleration may then decrease the sensitivity (Xu 2013). Event-related designs with shorter trial events may therefore benefit more from higher temporal resolutions, and this seems to be the case in the event-related faces/places task, though even here the gains seen in ROI measures are not impressive.

What these data demonstrate overall is that substantial reductions in TR do not produce strong benefits in statistical reliability in a straightforward manner. As has been shown previously (e.g. Smith et al., 2013) resting-state fMRI can substantially benefit from higher temporal resolution scanning, but this is not necessarily true for task paradigms, particularly when using ROI-based measures. One intriguing counter-example is the MVPA of the faces/places task data, which showed a largely consistent gain in sensitivity with the higher speed sequences; an advantage that produced statistically significant results in the multiband sequences, but not in the standard one. In this case, the larger number of data points may have served to produce more robust (i.e. less noisy) estimates of the average pattern of activity across trials, which in turn led to more reliable classifier performance. Much more work will be needed to substantiate this finding, and tease out the interaction between these novel acquisition schemes, and this also relatively novel analysis method.

Results from the two different scanner platforms are generally relatively symmetrical, with only minor differences. Scanner 2 produced a more coherent pattern of significant differences between the sequences in experiment 1, while Scanner 1 produced a (marginally) more coherent pattern in experiment 2. The two scanners tested have identical field strength (2.89T) and identical acquisition/reconstruction software was used on both, but Scanner 1 is a long, narrow-bore magnet and Scanner 2 is a short, wide-bore magnet. A number of hardware differences are important for the performance of multiband sequences. The design and size of the transmitting RF body coil differs, changing SAR and RF power requirements for the high multiband factor RF pulses. The gradient systems also differ, especially in their ability to actively cool from the high duty cycle EPI readouts. Disadvantages from the heating of the gradient system are manifold, but for multiband sequences the concomitant heating of the body coil changes the B1 field imparted. Finally, the main field (B0) homogeneity is more uniform and more stable in the long narrow-bore design. The relatively equivalent results on both systems are reassuring, in that researchers can be confident that results will likely be reasonably generalizable across other scanner platforms.

We sought to perform an evaluation of multiband acquisitions in a comprehensive and ‘real-world’ manner, using statistical outcome measures that working researchers tend to use. This entailed using a set of tasks chosen to give a range of effects within different experimental designs, and also collecting data from a set of human subjects. Much useful work in evaluating sequences can be done using MRI-phantoms and even simulation data, which certainly produce results with less variance. We sought to eliminate obvious subject-related confounds in our data by randomisation of the acquisition sequences within a scanning session, and randomising the order in which subjects took part in sessions on the two scanners. However, at least some of the variability in our results could be plausibly subject-related. As a measure of the ‘real-world’ performance of these sequences though, it could be argued that this is actually a more true reflection of their performance in such settings than phantom or simulation data would be. The tasks were chosen because they reliably produce well-replicated effects in a relatively short scan time, and they covered the two most common types of experimental design (block, and event-related). Clearly though, many other tasks also fit those criteria, and it is possible that there is some element in these particular tasks that confounded our results. Further testing with a greater range of paradigms and experimental designs would be ideal.

Based on these data, our recommendations for researchers interested in high-temporal resolution fMRI are essentially to proceed with caution. For resting-state fMRI there are obvious benefits, documented in the current data, and also by others (e.g. Smith et al., 2013). For task-based fMRI the picture is less clear, and any statistical benefit arising from a higher sampling rate is likely to depend on several factors, including the experimental design, the particular statistical outcome measure, and features of the analysis used.

## References

Auerbach, E. J., Xu, J., Yacoub, E., Moeller, S., & Uğurbil, K. (2013). Multiband accelerated spin-echo echo planar imaging with reduced peak RF power using time-shifted RF pulses. Magnetic Resonance in Medicine, 69(5), 1261–1267.

Barth, M., Breuer, F., Koopmans, P. J., Norris, D. G., & Poser, B. a. (2015). Simultaneous multislice (SMS) imaging techniques. Magnetic Resonance in Medicine, 81, n/a–n/a. doi:10.1002/mrm.25897

Beckmann, Mackay, Filippini, & Smith. (2009). Group comparison of resting-state FMRI data using multi-subject ICA and dual regression. NeuroImage, 47(Suppl 1), S148. doi:10.1073/pnas.0811879106

Bueti, D., & Walsh, V. (2009). The parietal cortex and the representation of time, space, number and other magnitudes. Philosophical Transactions of the Royal Society of London. Series B, Biological Sciences, 364(1525), 1831–1840. doi:10.1098/rstb.2009.0028

Boyacioğlu, R., Schulz, J., Koopmans, P. J., Barth, M., & Norris, D. G. (2015). Improved sensitivity and specificity for resting state and task fMRI with multiband multi-echo EPI compared to multi-echo EPI at 7T. NeuroImage, 119, 352–361.

Cauley, S. F., Polimeni, J. R., Bhat, H., Wald, L. L., & Setsompop, K. (2014). Interslice leakage artifact reduction technique for simultaneous multislice acquisitions. Magnetic resonance in medicine, 72(1), 93–102.

Chen, L., T. Vu, a., Xu, J., Moeller, S., Ugurbil, K., Yacoub, E., & Feinberg, D. a. (2015). Evaluation of highly accelerated simultaneous multi-slice EPI for fMRI. NeuroImage, 104, 452–459. doi:10.1016/j.neuroimage.2014.10.027

Epstein, R., & Kanwisher, N. (1998). A cortical representation of the local visual environment. Nature, 392(6676), 598–601.

Fias, W., Lammertyn, J., Reynvoet, B., Dupont, P., & Orban, G. a. (2003). Parietal representation of symbolic and nonsymbolic magnitude. Journal of Cognitive Neuroscience, 15(1), 47–56. doi:10.1162/089892903321107819

Fox, M. D., Snyder, A. Z., Vincent, J. L., Corbetta, M., Van Essen, D. C., & Raichle, M. E. (2005). The human brain is intrinsically organized into dynamic, anticorrelated functional networks. Proceedings of the National Academy of Sciences of the United States of America, 102(27), 9673–8. doi:10.1073/pnas.0504136102

Goulden, N., Khusnulina, A., Davis, N. J., Bracewell, R. M., Bokde, A. L., McNulty, J. P., & Mullins, P. G. (2014). The salience network is responsible for switching between the default mode network and the central executive network: replication from DCM. NeuroImage, 99, 180–90. doi:10.1016/j.neuroimage.2014.05.052

Hagberg, G. E., Zito, G., Patria, F., & Sanes, J. N. (2001). Improved detection of event-related functional MRI signals using probability functions. NeuroImage, 14(5), 1193–205. doi:10.1006/nimg.2001.0880

Jack, C. R., Bernstein, M. A., Fox, N. C., Thompson, P., Alexander, G., Harvey, D., … & Dale, A. M. (2008). The Alzheimer's disease neuroimaging initiative (ADNI): MRI methods. Journal of Magnetic Resonance Imaging,27(4), 685–691.

Ma, D. S., Correll, J., & Wittenbrink, B. (2015). The Chicago face database: A free stimulus set of faces and norming data. Behavior Research Methods, 47(4), 1122–1135.

Mitsis, G. D., Iannetti, G. D., Smart, T. S., Tracey, I., & Wise, R. G. (2008). Regions of interest analysis in pharmacological fMRI: how do the definition criteria influence the inferred result? NeuroImage, 40(1), 121–32. doi:10.1016/j.neuroimage.2007.11.026

Moeller, S., Auerbach, E., Van de Moortele, P. F., Adriany, G., & Ugurbil, K. (2008). fMRI with 16 fold reduction using multibanded multislice sampling. Proc Int Soc Mag Reson Med 16, 2366.

Moeller, S., Yacoub, E., Olman, C. A., Auerbach, E., Strupp, J., Harel, N., & Urbil, K. (2010). Multiband multislice GE-EPI at 7 tesla, with 16-fold acceleration using partial parallel imaging with application to high spatial and temporal whole-brain FMRI. Magnetic Resonance in Medicine, 63(5), 1144–1153. doi:10.1002/mrm.22361

Miller, K. L., Bartsch, A. J., Smith, S. M. (2015) Simultaneous multi-slice imaging for resting-state fMRI. Magnetom Flash, 63 (3), 70–77.

O’Craven, K. M., & Kanwisher, N. (2000). Mental imagery of faces and places activates corresponding stiimulus-specific brain regions. Journal of Cognitive Neuroscience, 12(6), 1013–23. doi:10.1162/08989290051137549

Pegors, T. K., Kable, J. W., Chatterjee, A., & Epstein, R. A. (2015). Common and unique representations in pFC for face and place attractiveness. Journal of Cognitive Neuroscience, 27(5), 959–73. doi:10.1162/jocn_a_00777

Peirce, J. W. (2007). PsychoPy—psychophysics software in Python. Journal of Neuroscience Methods, 162(1), 8–13.

Preibisch, C., Castrillón G., J. G., Bührer, M., & Riedl, V. (2015). Evaluation of multiband EPI acquisitions for resting state fMRI. PLoS ONE, 10(9), 1–14. doi:10.1371/journal.pone.0136961

Todd, N., Moeller, S., Auerbach, E. J., Yacoub, E., Flandin, G., & Weiskopf, N. (2016). Evaluation of 2D multiband EPI imaging for high-resolution, whole-brain, task-based fMRI studies at 3T: Sensitivity and slice leakage artifacts. NeuroImage, 124, 32–42. doi:10.1016/j.neuroimage.2015.08.056

Seeley, W. W., Menon, V., Schatzberg, A. F., Keller, J., Glover, G. H., Kenna, H., … Greicius, M. D. (2007). Dissociable intrinsic connectivity networks for salience processing and executive control. The Journal of Neuroscience 27(9), 2349–56. doi:10.1523/JNEUROSCI.5587-06.2007

Setsompop, K., Gagoski, B. A., Polimeni, J. R., Witzel, T., Wedeen, V. J., & Wald, L. L. (2012). Blipped-controlled aliasing in parallel imaging for simultaneous multislice echo planar imaging with reduced g-factor penalty. Magnetic Resonance in Medicine, 67(5), 1210–1224.

Shuman, M., & Kanwisher, N. (2004). Numerical magnitude in the human parietal lobe: Tests of representational generality and domain specificity. Neuron, 44(3), 557–569.

Smith, S. M., Fox, P. T., Miller, K. L., Glahn, D. C., Fox, P. M., Mackay, C. E., … & Beckmann, C. F. (2009). Correspondence of the brain's functional architecture during activation and rest. Proceedings of the National Academy of Sciences, 106(31), 13040–13045.

Smith, S. M., Beckmann, C. F., Andersson, J., Auerbach, E. J., Bijsterbosch, J., Douaud, G., … Glasser, M. F. (2013). Resting-state fMRI in the Human Connectome Project. NeuroImage 80, 144–168. doi:10.1016/j.neuroimage.2013.05.039

Xu, J., Moeller, S., Auerbach, E. J., Strupp, J., Smith, S. M., Feinberg, D. A., … & Uğurbil, K. (2013). Evaluation of slice accelerations using multiband echo planar imaging at 3T. NeuroImage, 83, 991–1001.

